# Breathlessness and the body: Neuroimaging evidence for the inferential leap

**DOI:** 10.1101/117408

**Authors:** Olivia K Faull, Anja Hayen, Kyle T S Pattinson

**Affiliations:** FMRIB Centre, University of Oxford, Oxford, UK; Nuffield Division of Anesthetics, Nuffield Department of Clinical Neurosciences, University of Oxford, Oxford, UK

**Author notes:** Corresponding author: Dr Olivia, Faull Nuffield Department of Clinical Neurosciences, University of Oxford, Oxford, UK.

**Keywords:** fMRI, breathlessness, symptoms, anxiety

## Abstract

Breathlessness debilitates millions of people with chronic illness. Mismatch between breathlessness severity and objective disease markers is common and poorly understood. Traditionally, sensory perception was conceptualised as a stimulus-response relationship, although this cannot explain how conditioned symptoms may occur in the absence of physiological signals from the lungs or airways. A Bayesian model is now proposed in which the brain generates sensations based on expectations learned from past experiences (priors), which are then checked against incoming afferent signals. In this model, psychological factors may act as moderators. They may either alter priors, or change the relative attention towards incoming sensory information, leading to more variable interpretation of an equivalent afferent input.

In the present study we conducted a preliminary test of this model in a supplementary analysis of previously published data (Hayen 2017). We hypothesised that individual differences in psychological traits (anxiety, depression, anxiety sensitivity) would correlate with the variability of subjective evaluation of equivalent breathlessness challenges. To better understand the resulting inferential leap in the brain, we explored whether these behavioural measures correlated with activity in areas governing either prior generation or sensory afferent input.

Behaviorally, anxiety sensitivity was found to positively correlate with each subject’s variability of intensity and unpleasantness during mild breathlessness, and with unpleasantness during strong breathlessness. In the brain, anxiety sensitivity was found to positively correlate with activity in the anterior insula during mild breathlessness, and negatively correlate with parietal sensorimotor areas during strong breathlessness.

Our findings suggest that anxiety sensitivity may reduce the robustness of this Bayesian sensory perception system, increasing the variability of breathlessness perception and possibly susceptibility to symptom misinterpretation. These preliminary findings in healthy individuals demonstrate how differences in psychological function influence the way we experience bodily sensations, which might direct us towards better understanding of symptom mismatch in clinical populations.

## Introduction

*“If the doors of perception were cleansed everything would appear to man as it is, infinite.*

*For man has closed himself up till he sees all things thro’ narrow chinks of his cavern*.” WILLIAM BLAKE, *The Marriage of Heaven and Hell*

The perception of bodily sensation is integral to the management of self within the environment. One frightening and debilitating perception is that of breathlessness, when breathing is perceived as inadequate and a threat to life. Breathlessness is experienced across a range of illnesses ^1,2^, including lung disease, heart disease and cancer. Breathlessness is notorious in that symptoms often are out of proportion to objective markers of disease ^3-7^. While perceptual systems have traditionally been considered to encompass a stimulus followed by the brain’s response, this relationship cannot explain the often-observed dissociation between perception and symptom extent, with extreme cases manifesting as medically unexplained symptoms ^8,9^. As it is the perception of symptoms that leads to their debilitating consequences, an overhaul is required in the way we consider the brain’s interaction with incoming sensory information. This would lead to better ways to understand and then treat unpleasant perceptions such as breathlessness.

With a launch into the Bayesian tidal wave of modern neuroscience ^10-15^, recent theories have proposed a comprehensive model of symptom perception ^16,17^. An important development of this model is the inclusion of a set of perceptual expectations, or ‘doors of perception’ in the words of William Blake. These perceptual ‘priors’ are neural representations of a distribution of expected values, which may be separated from the afferent neural inputs. Both priors and afferent sensory information can influence perception, which encompasses a range of probable perceptions (posterior distribution). Enhanced confidence in expectations (narrow, sharp priors) can increase their weight in the model, pulling the resulting perception away from the physiology and towards the prior. Furthermore, perceptual moderators exist within this system, such as anxiety ^18-20^, attention ^21-23^ or interoceptive ability ^24-27^, which may adjust either the prior expectations or incoming sensory information to influence perception. For instance, perception may be shifted to be higher or lower than the sensation, or there may be a greater range of possible perception values (widened distribution), which increases their ambiguity and susceptibility to misinterpretation and misclassification as a potential threat.

The ‘inferential leap’ to reconcile expectation and neural sensory information and form conscious perception occurs in the brain ^17^. One seductive theory consists of a division between agranular cortices (such as the anterior cingulate cortex and anterior insula) that generate prediction signals, and granular cortices (such as the primary sensory cortex and posterior insula), which compare afferent signals with predictions to generate prediction errors ^16,28,29^. It is hypothesized that behavioural factors such as decreased or redirected attention could also reduce the gain of sensory information within granular cortices ^30^, thereby diminishing the prediction error by increasing the relative weight of the priors in the model ^16,30^ Alternatively, behavioural influences may reduce the gain of the prior within agranular cortices ^16^ to reduce prediction errors and influence perception.

In this short report we have firstly investigated whether behavioural scores of anxiety, depression and anxiety sensitivity relate to the distribution of subjective scores (posterior perceptual distribution) of experimentally induced breathlessness. Mild and strong breathlessness were indicated by a conditioned stimulus (a shape presented on a screen), and implemented after a short anticipation period. Both levels of breathlessness were considered, as sensory afferents may be more vague or indefinite during mild breathlessness stimuli and might thus rely more heavily on priors. To do this we have undertaken a supplementary analysis on previously unreported aspects of a recently published study ^31^, to explore where in the brain these perceptual moderators act to alter perception.

## Materials and Methods

This study aimed to characterise functional brain activity during perception of a conditioned mild and strong breathlessness stimuli in 19 healthy participants (10 females, mean age ± SD, 24 ± 7 years). An account of conditioned responses to strong breathlessness has been published previously^31^, while the mild breathlessness stimulus was not considered due to its large between-subject variability. In the current report we have undertaken a more detailed evaluation of how behavioural measures relate to subjective stimulus perceptions in the mild condition, and where in the brain these perceptions may be modulated. Please see Hayen et al. (2017) ^31^ for a complete description of data acquisition and the lower level functional magnetic resonance imaging (fMRI) analysis. The study of Hayen et al. was a double-blinded placebo-controlled study of the effect of an opioid (remifentanil) on breathlessness, but in the present paper we are only considering the placebo condition (infusion of 0.9% saline).

### Behavioural questionnaires

The Center for Epidemiologic Studies Depression Scale (CES-D ^32^) was used to identify (and exclude) participants with clinical depression. The trait scale of the Spielberger State-Trait Anxiety Inventory (STAI ^33^) was used to characterize general participant anxiety. The Anxiety Sensitivity Index (ASI ^34^) was used to differentiate sensitivity to symptoms of anxiety in the form of bodily perceptions.

### Conditioned breathlessness and functional brain scanning

Scanning was conducted using a 3 Tesla Siemens Trio scanner, with physiological monitoring and control of end-tidal gases (see Hayen et al., 2017 ^31^). Briefly, an aversive delay-conditioning session was performed outside of the scanner, followed by two fMRI sessions on consecutive days (remifentanil or saline placebo, counterbalanced across participants). Participants learned associations between three visual cues and three respiratory sensations during the conditioning session, which were mild breathlessness, strong breathlessness or no breathlessness (unloaded breathing). The breathlessness stimulus used in this study was intermittent resistive inspiratory loading for 30 to 60 seconds, administered via an MRI compatible breathing system ^31^. Expiration was unrestricted via a one-way valve (Hans Rudolph, Shawnee, Kansas, USA). The stimuli were each presented four times during the scanning session in a semi-randomised, counterbalanced order, with a preceding anticipation period followed by a resistive loading stimulus (where appropriate). Immediately following each stimulus, participants were asked to rate both the intensity and unpleasantness of the preceding load on a visual analogue scale (VAS: 0-100%).

### Behavioural and fMRI analysis

In this short report we will only consider the fMRI session with the saline infusion. Full details on analysis procedures have been previously reported ^31^, and involved robust physiological noise correction of fMRI images. Whilst former analyses examined mean brain responses to anticipation and breathlessness (and the changes induced by remifentanil), the focus of this analysis was to explore how behavioural measures relate to the mean and variability of breathlessness perceptions in each subject, and to any corresponding changes in brain activity.

Mean and variability (standard deviation) of mouth pressure, subjective intensity and unpleasantness during scanning for both mild and strong loading were calculated for each subject. A full correlation matrix was then created on all behavioural and physiological variables, including questionnaires, mouth pressure and subjective breathlessness scores for each level of loading. As the behavioural variable of ASI score was shown to significantly correlate with trial-by-trial variation (standard deviation) of subjective scores, the group fMRI analysis previously reported ^31^ was adjusted to include a group mean and ASI score regressor. This analysis aimed to identify where functional brain activity correlates with differences in ASI score and thus extent of perceptual variability across subjects during saline administration, using whole-brain correction for multiple comparisons in FSL (FMRIB’s Software Library, www.fmrib.ox.ac.uk/fsl).

## Results

### Behavioural correlation matrix

Trait anxiety and depression were highly correlated across subjects, but neither correlated with ASI score (Figure 1). No behavioural scores (depression, trait anxiety or anxiety sensitivity) were found to significantly correlate with mean inspiratory pressure or subjective breathlessness VAS scores of intensity or unpleasantness for either mild or strong breathlessness conditions (Figure 1). When behavioural scores were compared to variability (standard deviation) in physiology and subjective scores, ASI was found to significantly correlate with variation in unpleasantness during both mild and strong breathlessness, and intensity with mild breathlessness (Figures 1 and 3). Both trait anxiety and depression were strongly correlated with the variation in pressure trace during strong (but not mild) breathlessness, but not subjective scores.

**Figure 1.**
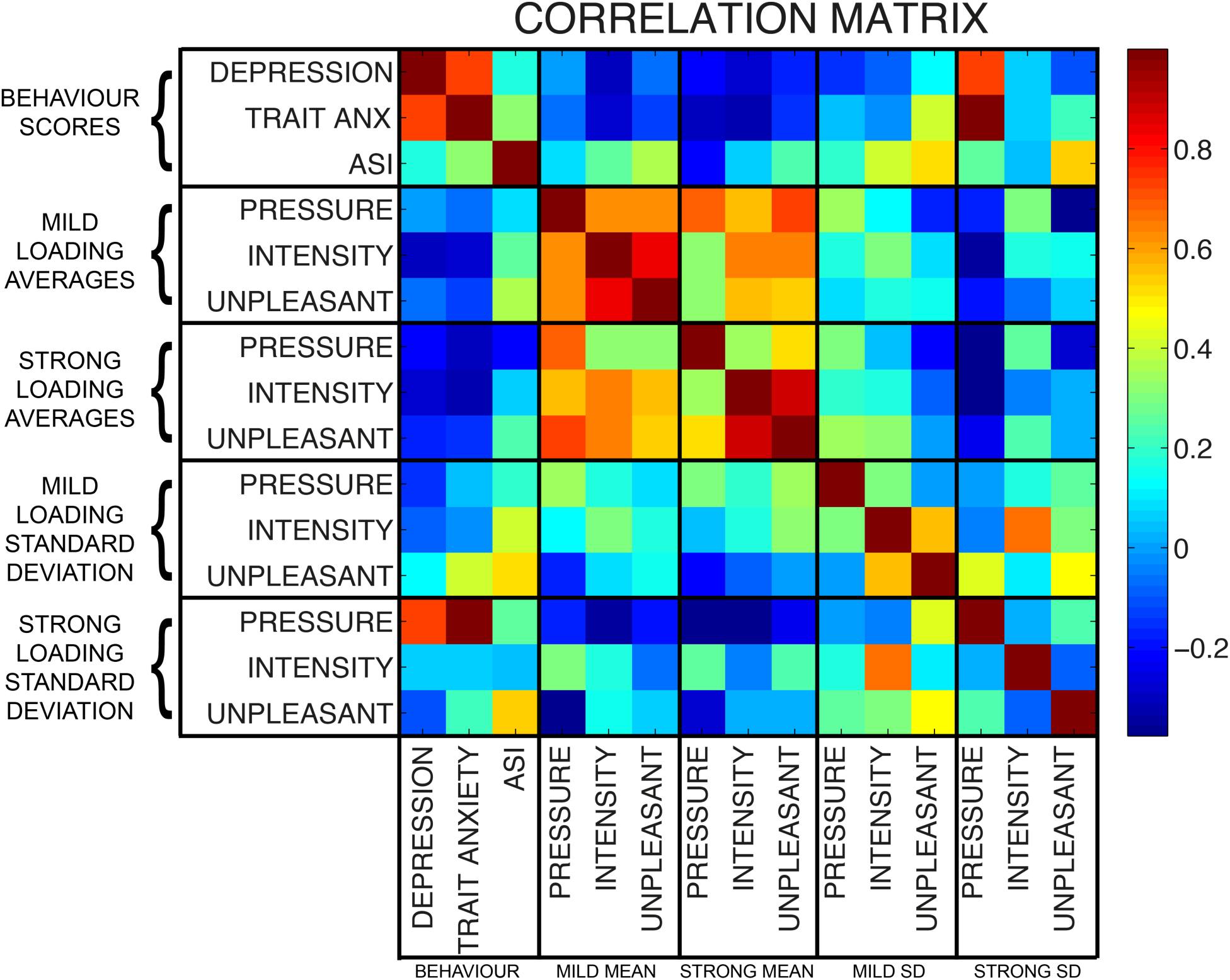
Full correlation matrix of all measured behavioural and physiological variables. Behavioural scores consisted of measures of depression, trait anxiety and anxiety sensitivity index (ASI). Mean and standard deviation measures of mouth pressure, intensity and unpleasantness scores are included for mild and strong resistive loading (breathlessness).

When mean subjective breathlessness scores and physiology were compared, average pressure, subjective intensity and unpleasantness were all strongly correlated during mild breathlessness (Figure 1). However, during strong breathlessness, intensity and unpleasantness scores became even more strongly correlated while ‘de-coupling’ from measures of inspiratory pressure. Lastly, while variation in intensity and unpleasantness scores were correlated during mild breathlessness, neither was reflective of variation in inspiratory pressure for either level of breathlessness.

### Average brain activity during anticipation and breathlessness

Conditioned associations between visual stimuli and breathlessness stimuli were confirmed prior to scanning in all subjects. Group mean brain activity during strong anticipation and breathlessness have been previously reported ^31^. No significant mean activity was observed during anticipation of mild breathlessness, and brain activity during mild and strong breathlessness is illustrated in Figure 2.

**Figure 2.**
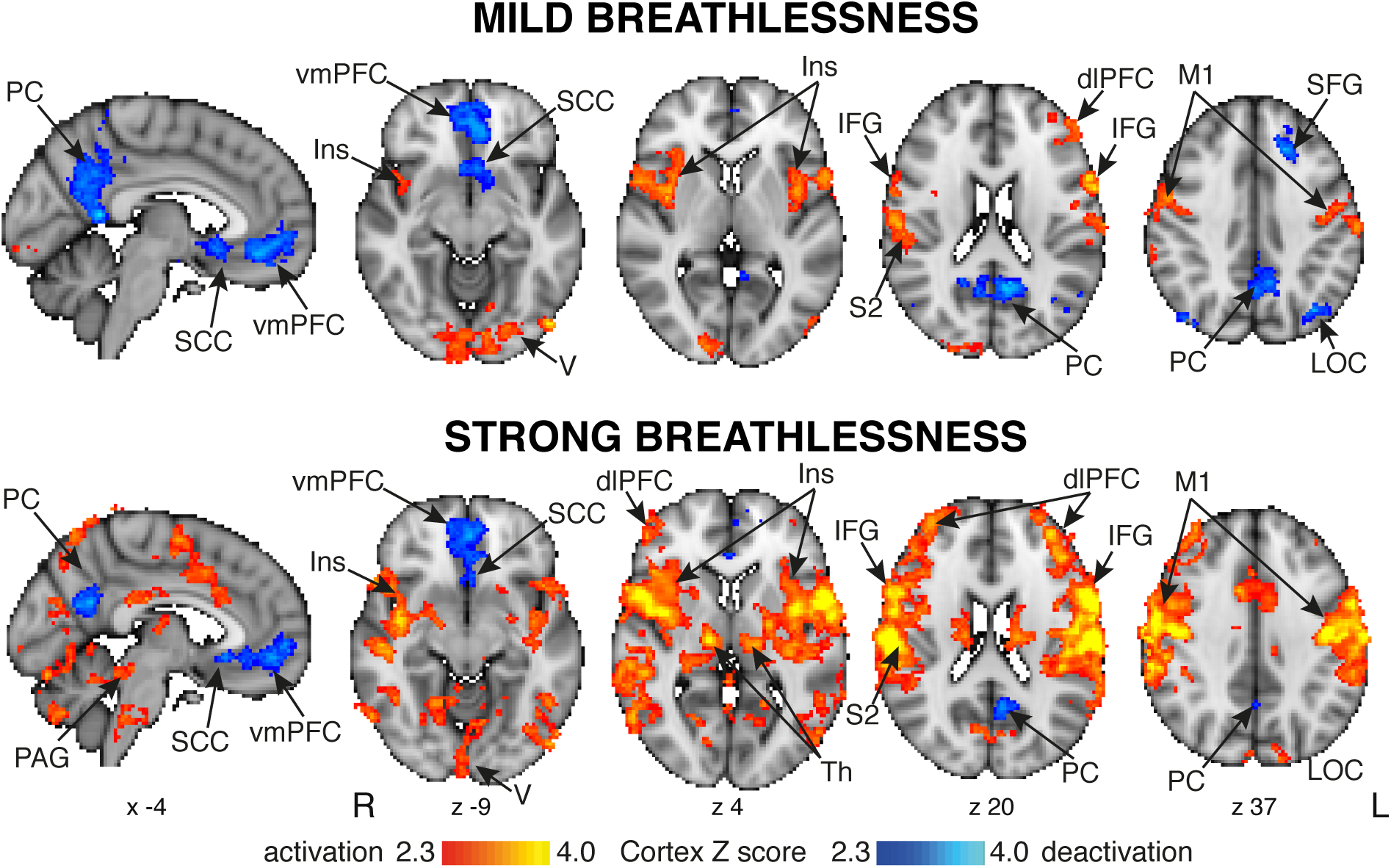
Mean BOLD changes identified during mild and strong breathlessness stimuli. The images consist of a colour-rendered statistical map superimposed on a standard (MNI 2×2×2 mm) brain. Significant regions are displayed with a threshold *Z* > 2.3, with a cluster probability threshold of *p* < 0.05 (corrected for multiple comparisons). Abbreviations: vmPFC, ventromedial prefrontal cortex; dlPFC, dorsolateral prefrontal cortex; SCC, subcingulate cortex; Ins, insula; IFG, inferior frontal gyrus; SFG, superior frontal gyrus; M1, primary motor cortex; S2, secondary somatosensory cortex; PC, precuneus; Th, thalamus; LOC, lateral occipital cortex; PAG, periaqueductal gray.

### Perceptual variation during mild breathlessness

During mild breathlessness, the extent of perceptual variation in subjective scores of both breathlessness intensity (r = 0.406, p = 0.048) and unpleasantness (r = 0.547, p = 0.010) were correlated with ASI score. When ASI score was subsequently investigated as a modulator of brain activity during mild breathlessness, it was found to correlate with brain activity in the left anterior insula only (Figure 3). No significant activity was found to correlate with ASI score during anticipation of mild breathlessness.

**Figure 3.**
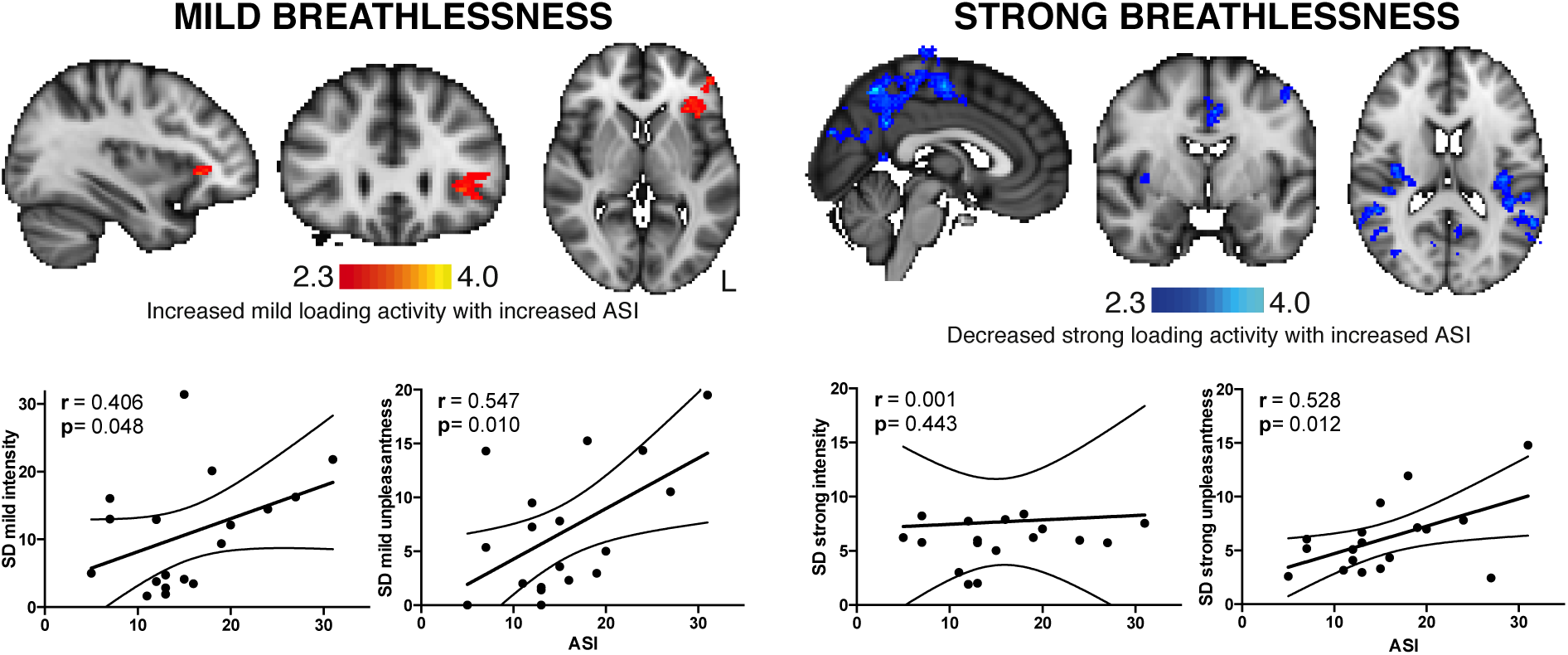
Relationship between perceptual variation, behavioural ASI score and brain activity. Left: (Top) Brain activity in the anterior insula that correlates with ASI score, and (bottom) significant correlations between ASI score and variation (standard deviation) in both intensity and unpleasantness during mild breathlessness. Right: (Top) Brain activity in the posterior insula, primary motor and sensory cortices, precuneus and posterior cingulate cortex that negatively correlates with ASI score, and (bottom) significant correlation between ASI score and unpleasantness, but not intensity during strong breathlessness.

### Perceptual variation during strong breathlessness

During strong breathlessness, the extent of perceptual variation in subjective scores of breathlessness unpleasantness was correlated with ASI score (r = 0.528, p = 0.012). Variation in breathlessness intensity no longer correlated with ASI score (r = 0.001, p = 0.443). ASI score was found to negatively correlate with activity in the posterior insula cortex, primary and secondary somatosensory cortices, primary motor cortex, dorsal anterior cingulate cortex, lateral occipital cortex and the precuneus cortex (Figure 3). No significant brain activity was found to correlate with ASI score during anticipation of strong breathlessness.

## Discussion

In this study we have shown that the greater an individual’s anxiety sensitivity index (ASI) score, the greater the variability in breathlessness scores to a set of standardised breathlessness challenges. Furthermore, during mild breathlessness, ASI score was found to correlate with brain activity in the anterior insula. Conversely, during strong breathlessness, ASI score was inversely correlated with activity in parietal primary sensorimotor cortices.

The extent of negative emotions such as anxiety and depression have long been considered potential modulators of perception ^18,19,21,23,25,35^. However, in healthy populations these scores may not be sensitive enough to identify a potential role in the interoceptive sensations of breathlessness. In contrast, anxiety sensitivity is a measure of alertness or sensitivity to bodily sensations of anxiety, and worry about the consequences of those sensations ^34^. Interestingly, in this report we have shown that it is an individual’s anxiety sensitivity that correlates with the extent of their variability in perceived breathlessness, and not generalized trait anxiety or depression. This attention and vigilance towards bodily sensations might thus render symptoms more ambiguous and susceptible to misinterpretation. Comparatively, trait anxiety and depression instead correlated with mouth pressure variability during strong breathlessness, indicating that participants with high trait anxiety might have modulated their breathing to avert negative sensations, and actively mediate the relationship between symptoms and expected perception.

Numerous previous studies have used a range of breathlessness stimuli to investigate where breathlessness symptoms are processed in the brain ^31,36-41^. What we have learned is that an extensive network of sensorimotor, affective and stimulus valuation areas are all highly active during breathlessness, as it is such a multi-dimensional experience ^6,7,42,43^. Moving forward, the challenge involves teasing apart where expectations (priors) and neural sensory information meet within this network to allow inference and perception. While studies using conditioned breathlessness cues can help us to understand the generation of priors ^44^, in this report we additionally investigated the perceptual variability around a repeated stimulus to probe how behavioural measures of anxiety, depression and anxiety sensitivity may be influencing the distribution of breathlessness scores, and where in the brain this may occur.

Within the Bayesian framework, the final perception of symptoms such as breathlessness is represented by a set of probable breathlessness values (posterior distribution). Psychological traits such as anxiety sensitivity could either interact with expectations, or with incoming sensory information to alter this posterior distribution ^17^. As this Bayesian system strives for efficiency, it aims to minimize the differences between prior expectations and afferent sensory information (prediction errors) ^28^. This could occur either by changing prior expectations, or reducing the importance (gain) of sensory neural information to lessen prediction errors. It has been elegantly hypothesized that aspects of this Bayesian framework may be somewhat anatomically distinct within the brain. Specifically, prior generation and predictions occur within the deep layers of agranular cortices such as anterior cingulate cortex and anterior insula 16,17,28,29, which are comprised of many projection neurons connected to granular cortices ^29,45-47^. Granular cortices, such as the primary sensory cortex and posterior insula, consist of well-differentiated layers including granule cells in layer IV that amplify thalamic sensory inputs ^48-50^.

In the current study, participants were conditioned to associate an abstract cue with upcoming mild or strong breathlessness. This learnt association allows the generation of breathlessness expectations, and we were then able to investigate where in the brain the behavioural anxiety sensitivity interacts with brain activity. During mild breathlessness, we observed a correlation with activity in the anterior insula (agranular), which has been previously implicated in prior generation within an interoceptive prediction system ^16^. Conversely, during strong breathlessness, anxiety sensitivity was inversely correlated with granular cortices such as the posterior insula and primary sensory cortex ^29,51,52^. Therefore, it is possible that anxiety sensitivity interacts within this Bayesian framework at either the level of the prior or at the level of receiving afferent inputs to the system, depending on the level of intensity of the stimulus. As anxiety sensitivity represents attention towards bodily sensations, it is possible that down-weighting priors and concentrating attention towards afferent sensation makes this system less robust, and as a result creates a wider posterior perceptual distribution. Comparatively, during stronger (and less ambiguous) breathlessness, anxiety sensitivity correlates only with perceptual variation of unpleasantness, but no longer intensity. The corresponding changes in granular cortex may represent modulation of the gain of afferent information, attempting to bring sensations closer to priors to reduce prediction errors.

### Clinical Relevance

The current study has been carried out in healthy volunteers with no history of respiratory disease. Studying healthy populations can aid us in understanding normal variants in physiology, psychology and perception. Still, the challenge remains to apply these concepts to clinical populations. If an individual suffers from chronic breathlessness, they may (over time) alter their priors and thus change their perception.

This may result in a shift of the prior further from the neural sensory information (a leftward or rightward shift of the prior illustration in Figure 4). It remains to be investigated how this change in expectation within the course of chronic disease may be influenced by pre-existing behavioural levels of anxiety, depression and anxiety sensitivity. This could help to explain how treatment options such as pulmonary rehabilitation for chronic obstructive pulmonary disease (COPD) may be addressing these expectations of breathlessness ^53^ and determine in which populations and under what conditions such measures would be expected to work best. Using the Bayesian framework to link relevant baseline measures of anxiety and interoceptive sensitivity to neural activation within clinical populations could also help to understand and address maladaptive perceptual differences, e.g. dangerous ‘under-’ and ‘over-’ perception of symptoms in asthma sufferers.

**Figure 4.**
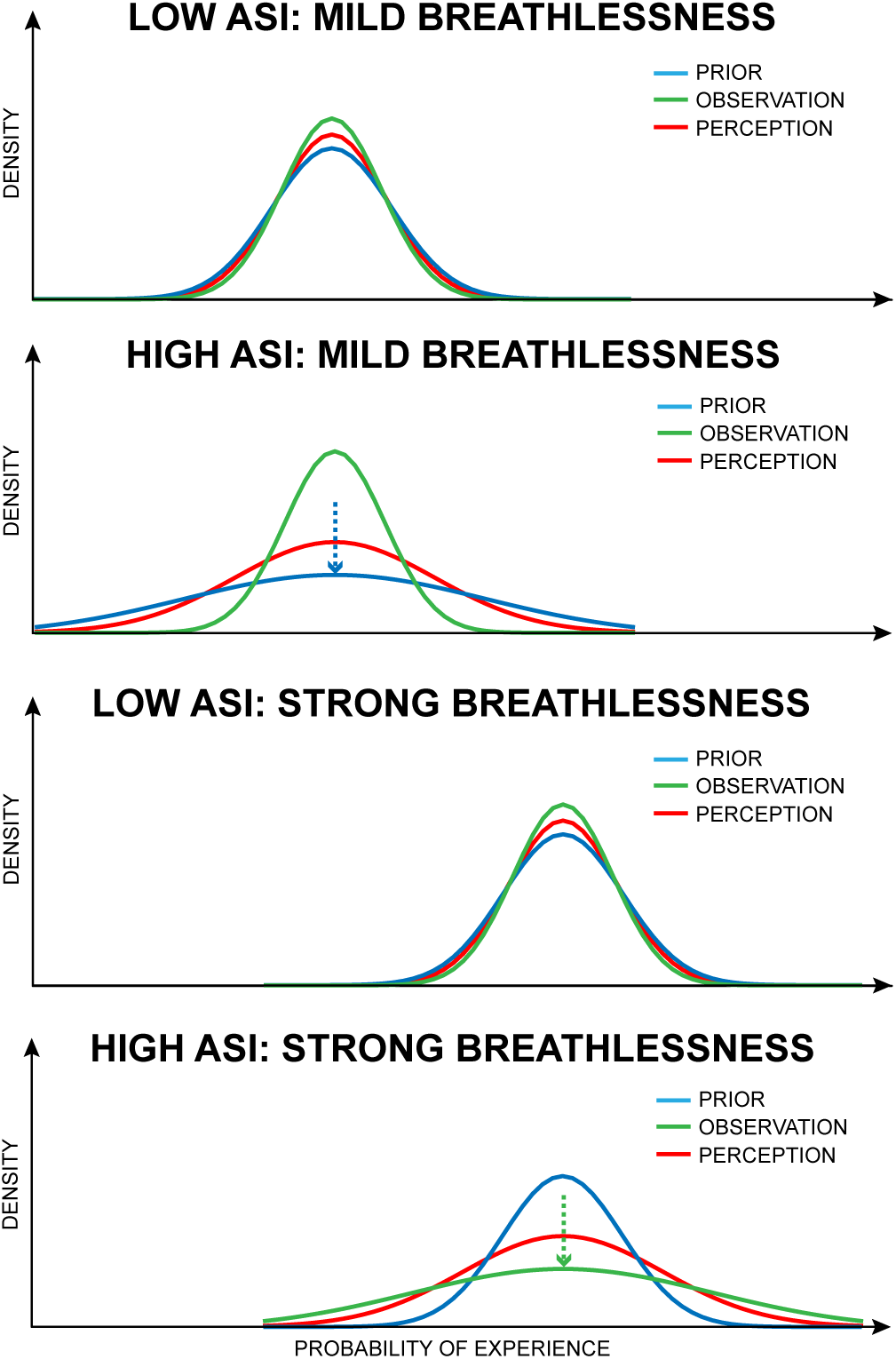
Theoretical possible relationships between ASI and posterior distribution of breathlessness perception using a Bayesian framework. Top two panels: Minimal influence of priors and sensory input (observation) on posterior distribution with low ASI, but flattened prior may widen posterior perceptual distribution with high ASI during mild breathlessness. Bottom two panels: Minimal influence of priors and sensory input (observation) on posterior distribution with low ASI, but flattened observation may widen posterior perceptual distribution with high ASI during strong breathlessness. Figure adapted from Van den Bergh et al. (2017).

### Limitations

This study is a supplementary analysis of previously published work, representing preliminary pilot data in healthy volunteers with small study numbers (n = 19) and limited stimulus repetitions (n = 4 each for mild and strong breathlessness). Whilst previously published research has demonstrated both improved ^54^ and worsened ^55^ respiratory perceptual accuracy with greater anxiety, the current results showed no effect of trait anxiety on perception. Rather, we have observed a relationship between anxiety sensitivity and perceptual variation. While anxiety sensitivity represents a separate facet of anxiety constrained to bodily sensations ^34^, numerous other variables may also contribute to differences with previously published results. These factors may include the continuous ratings used in this study compared to categorical ratings used previously ^56^, the relatively low trait anxiety values of the study subjects (mean 34 ± 9 (SD), compared to previous classifications of low (29) and high (55) trait anxiety ^57^), and/or the small subject numbers and repeats employed.

This study was also unable to determine the location and shape of the prior in relation to both the sensory observation and resulting perceptual (posterior) distribution.

It is possible that anxiety sensitivity, anxiety and / or depression induce a lateral shift of the prior, and our assumed changes in prior shape are inferred from the resulting changes in perceptual variation. It is clear that further work is required to explore the relationship between anxiety sensitivity and prior generation, and how this may change across a broad spectrum of generalized anxiety, to determine its place within the Bayesian symptom perception framework.

## Conclusions and future directions

This short report is a preliminary insight into potential mechanisms of perceptual modulation of breathlessness within the Bayesian framework. Within this framework, the brain integrates prior expectations with afferent sensory information to create breathlessness perception. Behavioral modulators could potentially alter this relationship and influence subsequent perceptual distributions. Here, we have shown that level of anxiety sensitivity explains variations in breathlessness perception between healthy volunteers, possibly modifying both priors and afferent sensations, which are processed in distinct brain areas. Therefore, attention to bodily sensations (ASI) may reduce the robustness of this system in healthy individuals, and increase susceptibility to misinterpretation of breathlessness. Future work on larger cohorts needs to address the relationship between anxiety sensitivity, interoceptive accuracy/confidence and breathlessness perceptions, to investigate how both attention to bodily sensations and interoceptive abilities may interact to adjust the doors of symptom perception.

**Table 1.** Effects of loading on respiratory parameters. P_ET_CO_2_=partial pressure of end-tidal carbon dioxide. P_ET_O_2_=partial pressure of end-tidal oxygen. Values are presented as mean (SD). N=19. Complete heart rate data in each epoch only available for 15 subjects. * = significantly different from saline unloaded breathing at p<0.001. ¶ = significantly different from saline anticipation unloaded breathing at p<0.05

| Variable | Anticipation unloaded | Unloaded breathing | Anticipation mild | Mild loading | Anticipation strong | Strong loading |
| --- | --- | --- | --- | --- | --- | --- |
| Mouth pressure amplitude [cmH<sub>2</sub>O] | 2.7 (0.7) | 2.4 (0.5) | 2.6 (0.7) | 4.0 (0.8) | 3.5 (1.7)¶ | 12.7 (4.1)* |
| $P_{ET}CO_2$ [kPa] | 5.5 (0.6) | 5.6 (0.6) | 5.6 (0.5) | 5.5 (0.5) | 5.5 (0.5) | 5.5 (0.6) |
| $P_{ET}O_2$ [kPa] | 20.0 (0.9) | 19.8 (0.8) | 19.8 (0.7) | 20.2 (0.9) | 19.9 (0.7) | 20.2 (0.8) |
| Intensity rating [%VAS] |  | 12 (16) |  | 32 (21) |  | 71 (20)* |
| Unpleasantness rating [%VAS] |  | 10 (18) |  | 25 (25) |  | 61 (32)* |
| Heart rate [min<sup>-1</sup>] (N=15) | 68 (11) | 67 (10) | 69 (9) | 67 (12) | 68 (11) | 69 (11) |

## Acknowledgements

This research was supported by the Medical Research Council (UK). This research was further supported by the JABBS Foundation and the National Institute for Health Research, Oxford Biomedical Research Centre based at Oxford University Hospitals NHS Trust and University of Oxford. The authors would like to thank Omer Van Den Bergh for his thoughts and comments on the manuscript.

